# Cardiac unloading via a transvalvular pump prevents arrhythmias following myocardial ischemia / reperfusion by preserving coronary microcirculation and cardiomyocyte nanostructure

**DOI:** 10.64898/2026.10.14.758560

**Authors:** Madison Ammon, Pedro H. C. Melo, Kacper Nowak, Gerard Nuovo, Alexandre Abizaid, Juan F. Granada, Daniel Burkhoff, Grzegorz L. Kaluza, Alfred A. Kyrollos, Jerald Curran, Sathya Dev Unudurthi, Rengasayee Veeraraghavan

## Abstract

Cardiac damage in the aftermath of myocardial ischemia / reperfusion (MI/R) compromises both electrical and mechanical function, predisposing patients to arrhythmia and heart failure. Microvascular obstruction (MVO) is increasingly recognized as a sequela of MI/R and a contributor to proarrhythmic cardiac remodeling. While transvalvular unloading (TVU) using a percutaneously delivered pump is garnering appreciation as a means of limiting infarct size and preserving mechanical function following percutaneous coronary intervention, its benefits for arrhythmia prevention remain unclear. We used a porcine model to test the hypothesis that cardiac unloading prior to and through reperfusion mitigates MVO, preserves cardiomyocyte nanostructure, and alleviates arrhythmias following MI/R. Anesthetized domestic swine undergoing acute ischemia (LAD occlusion, 90 mins) and reperfusion (120 mins) were randomized to TVU (n=5) and no treatment (control; n=6) treatments. Thioflavin-S was infused into the mid-LAD to assess coronary microcirculation and quantify MVO. Arrhythmia burden was assessed by *in vivo* electrocardiography using 3 lead EKG. Cardiac tissues were preserved for histology and optical microscopy studies. Compared to no treatment controls, TVU hearts evidenced significantly reduced infarct area (48 ± 5 % vs. 59 ± 11 % p = 0.03) and MVO (19 ± 4 % vs. 27 ± 5%, p = 0.03) normalized to area at risk and a higher myocardial salvage index (52 ± 5 % vs. 41 ± 11% p = 0.03). Extensive perivascular edema, and neutrophil infiltration observed in controls was markedly reduced in TVU hearts. Confocal microscopy and spatial pattern analysis revealed significantly mitigated disruption of gap junctions and Na_V_1.5 dissociation from N-cadherin in TVU hearts compared to controls. Importantly, severity-adjusted arrhythmia burden during reperfusion was significantly lower in TV-P hearts (8±8 vs. 12±5 episodes lasting greater than 5seconds in controls). Our results demonstrate that TVU effectively prevents arrhythmias during reperfusion by preserving coronary microvasculature and cardiomyocyte nanostructure.

## INTRODUCTION

Each year, as many as 300,000 patients undergo percutaneous coronary intervention (PCI) in the United States as a treatment for acute myocardial infarction (AMI)^1,2^. Current clinical practice prioritizes rapid restoration of blood flow to ischemic myocardium with the goal of minimizing infarct size and improving patient outcomes over both acute and chronic time frames^3,4^. However, the sudden restoration of blood flow to the ischemic territory is well-known to lead to ischemia-reperfusion injury (IRI). This process exacerbates myocardial injury though oxidative stress, inflammation, mitochondrial dysfunction and cell death and IRI^5^ contributes to increased arrhythmia burden, risk of heart failure, and other adverse outcomes in patients. Thus, there is a pressing need for effective therapies to ameliorate IRI.

Mechanical unloading of the heart prior to reperfusion has emerged as a potentially attractive therapeutic option. In pre-clinical studies, mechanically unloading the left ventricle reduces pressure and volume, decreases wall stress and myocardial oxygen consumption, enhances coronary perfusion, and activates cardioprotective signaling^6–11^. The Impella CP™ (Johnson & Johnson MedTech Heart Recovery, MA, USA) a percutaneously delivered transvalvular microaxial flow pump provides a clinically deployable means of mechanically unloading the heart prior to PCI. Impella actively unloads the left ventricle by continuously aspirating blood from the left ventricle into the ascending aorta. This reduces LV pressure, volume and cardiac work, and alleviates the stress placed on the heart prior to and during reperfusion^6,7,9–11^. Mounting evidence suggests that mechanically unloading the heart before reperfusion can reduce infarct size thereby decreasing the subsequent risk for developing heart failure^12,13^. However, the effect of mechanical unloading prior to PCI on arrhythmia burden following reperfusion remains unclear. Using a porcine model of acute myocardial ischemia / reperfusion (MI/R), we tested the hypothesis that transvalvular unloading (TVU) applied prior to reperfusion can mitigate proarrhythmic sequela of IRI and alleviate arrhythmia burden. By correlating quantitative results from multi-modal, multi-resolution imaging with electrocardiography, we demonstrate that TVU mitigates cardiac injury and inflammation, preserves myocardium and microcirculation, prevents proarrhythmic remodeling of cardiac nanostructure, and ultimately ameliorates ventricular tachycardia (VT) following AMI.

## METHODS

### Induction of Myocardial Infarction and Mechanical Circulatory Support

The study was approved by the Institutional Animal Care and Use Committee of the Cardiovascular Research Foundation (CRF) Skirball Center of Innovation, Orangeburg, NY. All animals received standard care outlined by the Guide to Care and Use of Laboratory Animals by the National Academy of Science and Animal Welfare Regulations.

A total of 11 domestic swine (43-52 kg) were anesthetized with a rapid-acting general intramuscular anesthetic (tiletamine + zolazepam, 2-5 mg/kg) supplemented by glycopyrrolate (0.005-0.02 mg/kg). The animals underwent endotracheal intubation and were maintained with continuous inhalation of 1-3% isoflurane. Arterial left and right carotid accesses were obtained, under general anesthesia, using either a cutdown or percutaneous technique.

Anticoagulation with heparin was achieved (3,000-10,000 U) to maintain activated coagulation time ≥300 seconds. The left anterior descending artery (LAD) was accessed with a 7F JL 3.5 guide catheter, and a coronary angioplasty balloon was positioned below the origin of the first diagonal branch. The balloon was inflated to achieve a balloon-to-artery diameter ratio of 1.2:1 (using quantitative coronary angiography – QCA), and total coronary occlusion was confirmed every 30 minutes through serial angiographies during a 90-minute balloon-occlusion period. The balloon was deflated in the 90-minute in all animals.

After the first 60 minutes of occlusion, swine were randomized to the transvalvular mechanical unloading group (TVU, n=5), or the control group (n=6). Control subjects had continued LAD occlusion alone for 30 minutes (n=6). TVU subjects had continued LAD occlusion for 30 minutes with concomitant activation of an Impella CP (n=5). After a total of 90min of LAD occlusion, both groups were subjected to 120minutes reperfusion. In the TVU group, Impella CP remained active throughout 120min of reperfusion. The Impella CP was inserted via a 14F sheath in the right carotid artery and activated at maximal support without suction. Limb-lead ECGs were collected throughout the duration of the experiment.

### Euthanasia and Postmortem Studies

At the end of the 120-minute reperfusion period, the JL 3.5 guide catheter was reintroduced to cannulate the left coronary, followed by a final angiography. Subsequently, an over-the-wire balloon was utilized to briefly reocclude the LAD. Through the lumen of this inflated ballon, 30 mL of a 4% thioflavin-S (T-S) solution was infused into the mid-LAD at the same site (immediately distal to the first diagonal of the left anterior descending coronary artery) used for the initial occlusion, enabling myocardial staining with T-S. T-S is a fluorescent dye that stains the endothelium of blood vessels that are being perfused and is a standard technique for identifying anatomic zones of no-reflow or microvascular obstruction in cardiac, cerebral, renal, and other tissue. When viewed under UV light, perfused tissue can be differentiated from non-perfused tissue by the presence or absence of fluorescence, respectively. After one minute, the balloon was removed, allowing T-S to circulate. Euthanasia was then performed using a solution containing pentobarbital sodium and phenytoin sodium.

Upon excision of the hearts, the left ventricle (LV) was sectioned into 10 mm thick short-axis slices from the apex to the basal level. Each slice was examined under ultraviolet (UV) light and photographed (apical view) with a reference ruler. The injection of T-S just before termination in this case served to directly visualize both the ischemic risk zone (area at risk) and the microvascular obstruction (MVO area). 8mm biopsy punches were collected for histological analysis and immunohistochemistry from the basal side of LV slices centered on area at risk as determined using S-thioflavin fluorescence under UV light. The area underlining each biopsy punch was stained using tissue dye (e.g., India ink) to ensure basal/apical orientation and fixed in 10% neutral buffered formalin for histological analysis. Next, LV rings were incubated in a warm 2% 2,3,5-triphenyl tetrazolium chloride (TTC) solution for 20 minutes and photographed under ambient light (apical view) for determination of infarct area.

### Histology

Tissues samples derived from our pig studies were fixed in formalin, embedded in paraffin, and prepared for histological studies as previously described^14,15^. Hematoxylin and eosin (H&E) staining of tissue sections was performed to visualize cells, cellular nuclei, and extracellular spaces. Neutrophils were identified via immunohistochemical staining of elastase as a measure of inflammation.

Additionally, ventricular tissue sections were obtained from infarct and remote myocardium (≥ 1cm from the infarct border) from pathology samples derived from human patients with acute MI (death occurred ≤ 72 hours following AMI) and immunohistochemistry was performed as previously described^14,15^ to detect the cardiac sodium channel, Na_V_1.5, and the cell-cell junction protein, N-cadherin (N-cad). In brief, the immunohistochemistry protocol used the Leica Bond Max automated platform along with the Fast red (DS 9820) and DAB (DS 9800) detection kits (Leica Biosystems, Buffalo Grove, IL). Optimal labeling conditions for each antibody were determined by testing various dilutions and pretreatment conditions. Co-expression analyses were performed using the Nuance / InForm system, whereby each chromogenic signal was separated, and overlap assessed to determine the extent of colocalization between the two proteins of interest as previously described^16–18^.

### Confocal Immunofluorescence

Immunofluorescent labeling of cryosections (5 µm thickness) of fresh-frozen myocardium was performed, as previously described^19,20^. Briefly, cryosections were fixed with paraformaldehyde (2%, 5 min at room temperature), permeabilized with Triton X-100 (0.2% in PBS for 15 min at room temperature) and treated with blocking agent (5% NDS, 0.1% triton in PBS for 2 h at room temperature) prior to labeling with primary antibodies (overnight at 4 °C). Samples were then washed in PBS (3 × 5 min in PBS at room temperature) prior to labeling with secondary antibodies. Samples were then labeled with goat anti-mouse and goat anti-rabbit secondary antibodies conjugated to Alexa 594 and STAR 635P were used (1:8000; Abberior). Samples were then washed in PBS (3 × 5 min in PBS at room temperature) and optically cleared using Scale U2 buffer (48 h at 4 °C) before being mounted in ProLong Glass (Invitrogen, Rockford, IL)^19,20^.

Samples were imaged using a TCS SP8 laser scanning confocal microscope and equipped with a pulsed supercontinuum laser (470–670 nm; 80 MHz pulse rate), a Plan Apochromat 63×/1.4 numerical aperture oil immersion objective, and liquid-cooled hybrid detectors with single photon detection capability (Leica, Buffalo Grove, IL). Time gating of light collection (1.5–3.5 ns following each laser pulse) was applied to exploit differences in fluorescence lifetime, and thereby, separate fluorescence emissions of Thioflavin-S dye from emissions of Alexa 594 and Star 635P. Images were collected as Nyquist-sampled z-stacks with fluorophores imaged sequentially. Images were analyzed using our previously validated point pattern analysis-based spatial analysis approach, Spatial Pattern Analysis using Closest Events (SPACE)^19,21,22^.

### Analysis of Electrocardiograms

Limb lead ECGs collected from our porcine studies were visualized in MATLAB and analyzed manually. Ventricular tachycardias were identified as three or more consecutive premature beats and durations of individual arrhythmic episodes tabulated to quantify ventricular tachycardia (VT) burden.

### Statistical Analysis

All statistical analyses were performed using R (R Foundation for Statistical Computing, Vienna, Austria). Histological variables (area at risk, infarct area, MVO, and myocardial salvage index) were expressed as mean ± SD and were compared between the TVU and control groups using the nonparametric Mann–Whitney U test. Given the directional hypothesis that TVU would reduce infarct size and mitigate MVO, P values for these histological comparisons were calculated as 1-sided, and values <0.05 were considered statistically significant.

## RESULTS

### TVU Mitigates Myocardial Damage, Microvascular Obstruction Following IRI

We examined the overall impact of mechanical unloading on severity of myocardial injury following MI/R. We performed histological studies to compare the myocardial area at risk and myocardial salvage index between groups (Fig.2). While the mean area at risk normalized to total LV area was similar between the transvalvular unloading (TVU) and control groups (46 ± 7 % vs. 42 ± 9 % respectively; p = 0.2), the infarct area normalized to total area at risk was significantly lower (48 ± 5 % vs.59 ± 11 % p = 0.03) and the myocardial salvage index was correspondingly higher (52 ± 5 % vs. 41 ± 11% p = 0.03) in TVU hearts. In addition, UV imaging of thioflavin-S dye perfused into hearts after MI/R revealed that TVU hearts had significantly less microvascular obstruction (MVO; normalized to area at risk; Fig. 2) relative to controls (19 ± 4 % vs. 27 ± 5%, p = 0.03).

**Figure 1:**
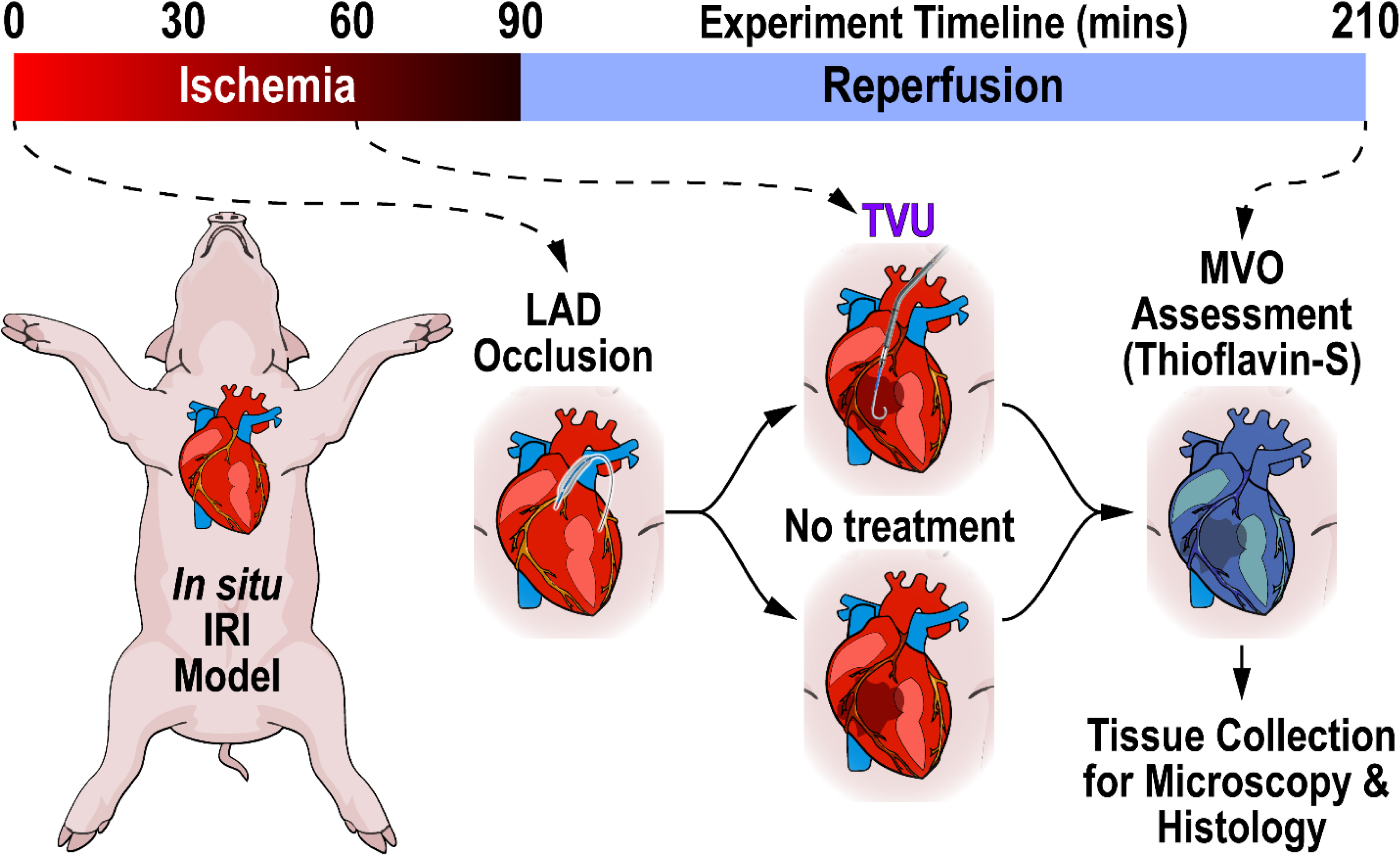
Schematic of the experimental protocol for porcine MI/R studies. Pigs underwent 90 minutes of ischemia and 120 minutes of reperfusion. Animals in the treatment group received TVU at 60 minutes into the ischemic period. Hearts were then perfused with a fluorescent marker of microcirculation (Thioflavin-S) and tissue preserved for microscopy and histology studies.

**Figure 2:**
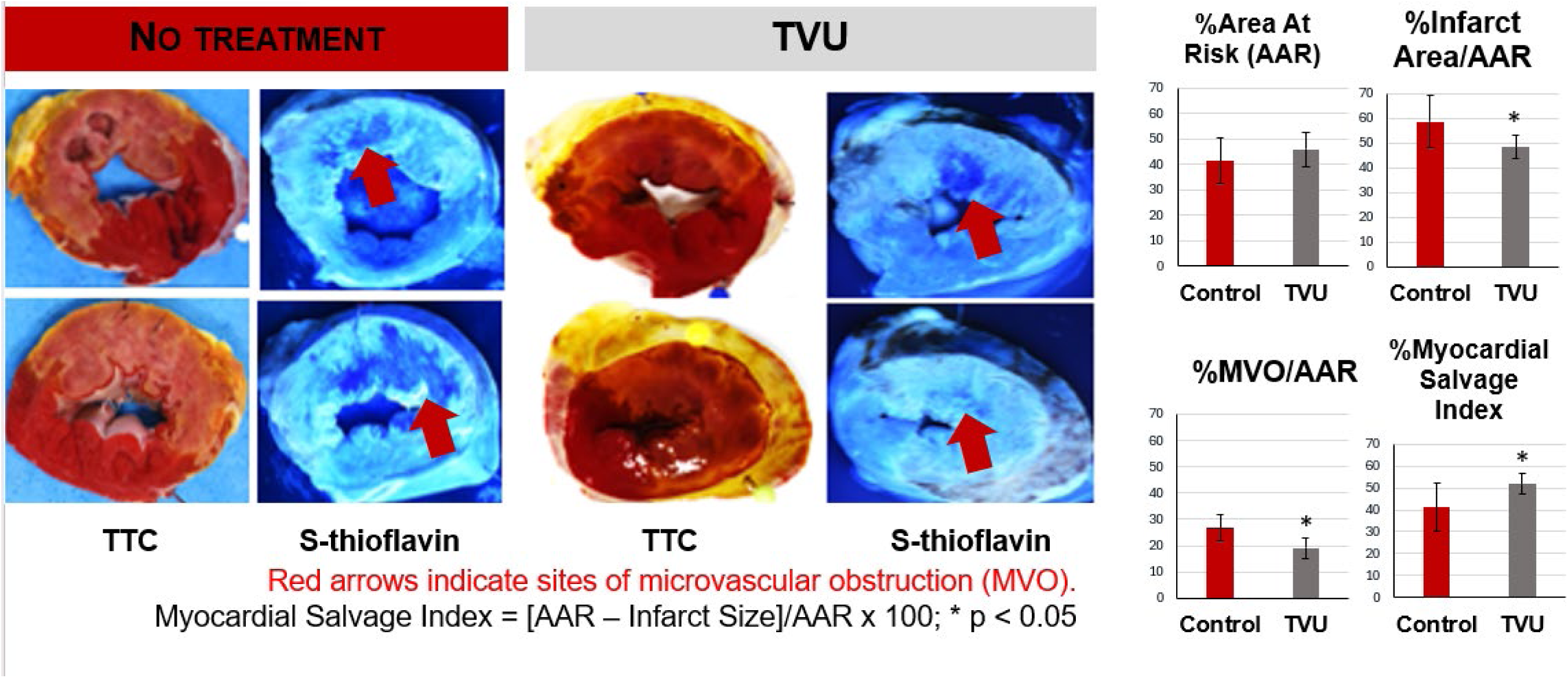
TVU reduces MVO and Infarct area. **A)** Representative images and **B)** summary quantification of myocardial tissue injury (TTC) and MVO (Thioflavin-S). * p <0.05 by Mann-Whitney U Test (One-Tailed) n = 6 for the no treatment group, n = 5 for the TVU group.

### TVU Mitigates Inflammation, Preserves Microvasculature, and Limits Cardiac Edema Following MI/R

Given the established association between microvascular dysfunction and inflammation in the context of cardiac arrhythmias^23–26^. we undertook histological investigation of the immune response and vascular barrier function. Neutrophil infiltration, quantified as the number of elastase-positive cells per cm^2^, was significantly reduced within the area at risk in TVU hearts relative to controls (12.9 ± 15 vs. 51.2 ± 29, p < 0.01). As expected, infiltration in AAR was significantly elevated relative to remote myocardium in TVU hearts (1 ± 0.4, p = 0.02 vs. area at risk) in TVU hearts; Fig. 4). Histological assessment showed a significantly lower degree of pericapillary in TVU hearts compared to control hearts following MI/R (Fig. 3).

**Figure 3:**
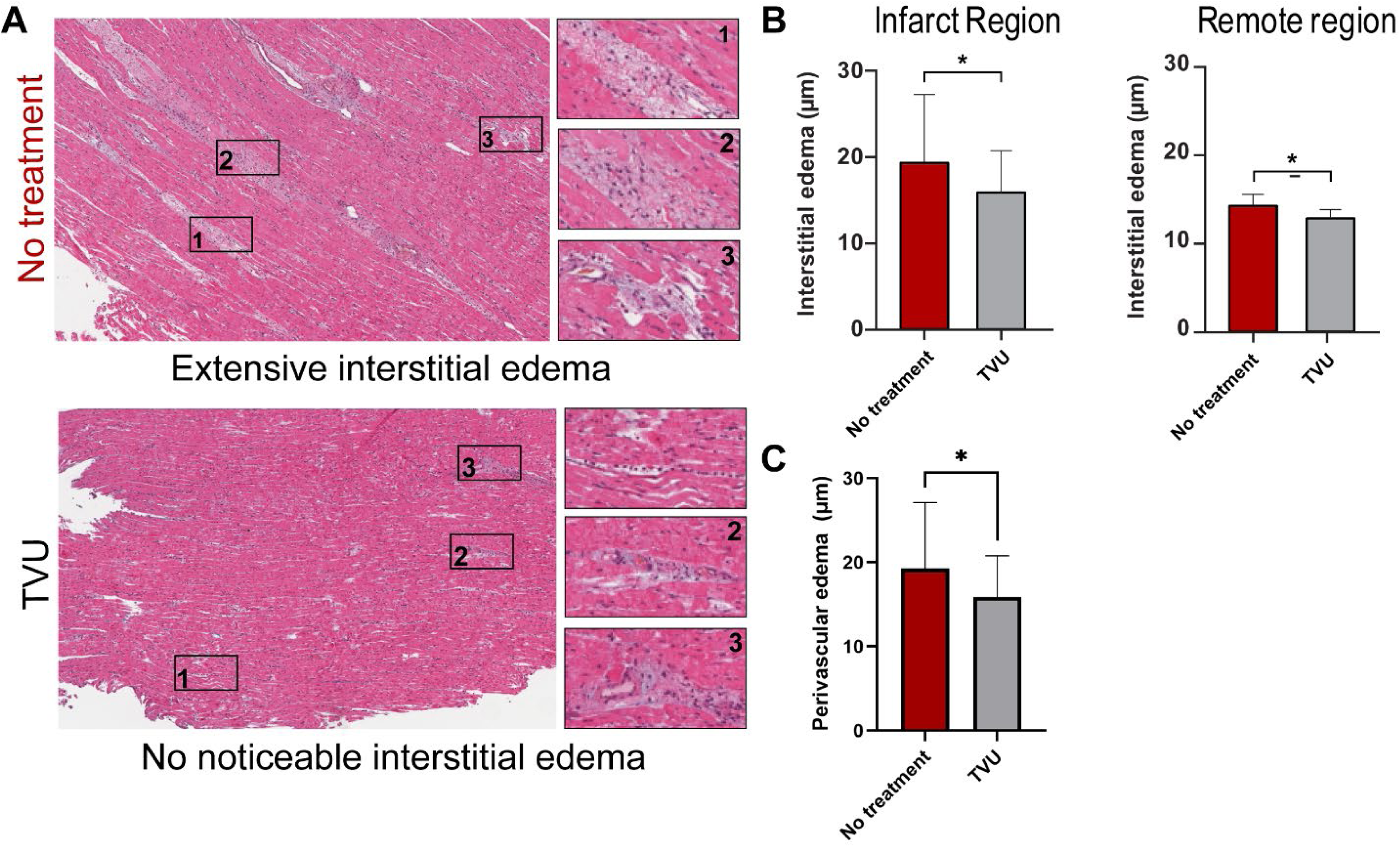
Histological observations of edema. **A)** Representative images (from infarct regions) and summary results demonstrating mitigation of **B)** interstitial edema and **C)** perivascular edema by TVU. Edema was quantified in terms of intercellular spacing. Perivascular sites (black boxes) evidence extensive edema in control (no treatment) hearts but not in TVU hearts. n = 3 pigs for no treatment and 5 for TVU; 12 regions of interest evaluated per heart; * p < 0.05 by Mann-Whitney test.

**Figure 4:**
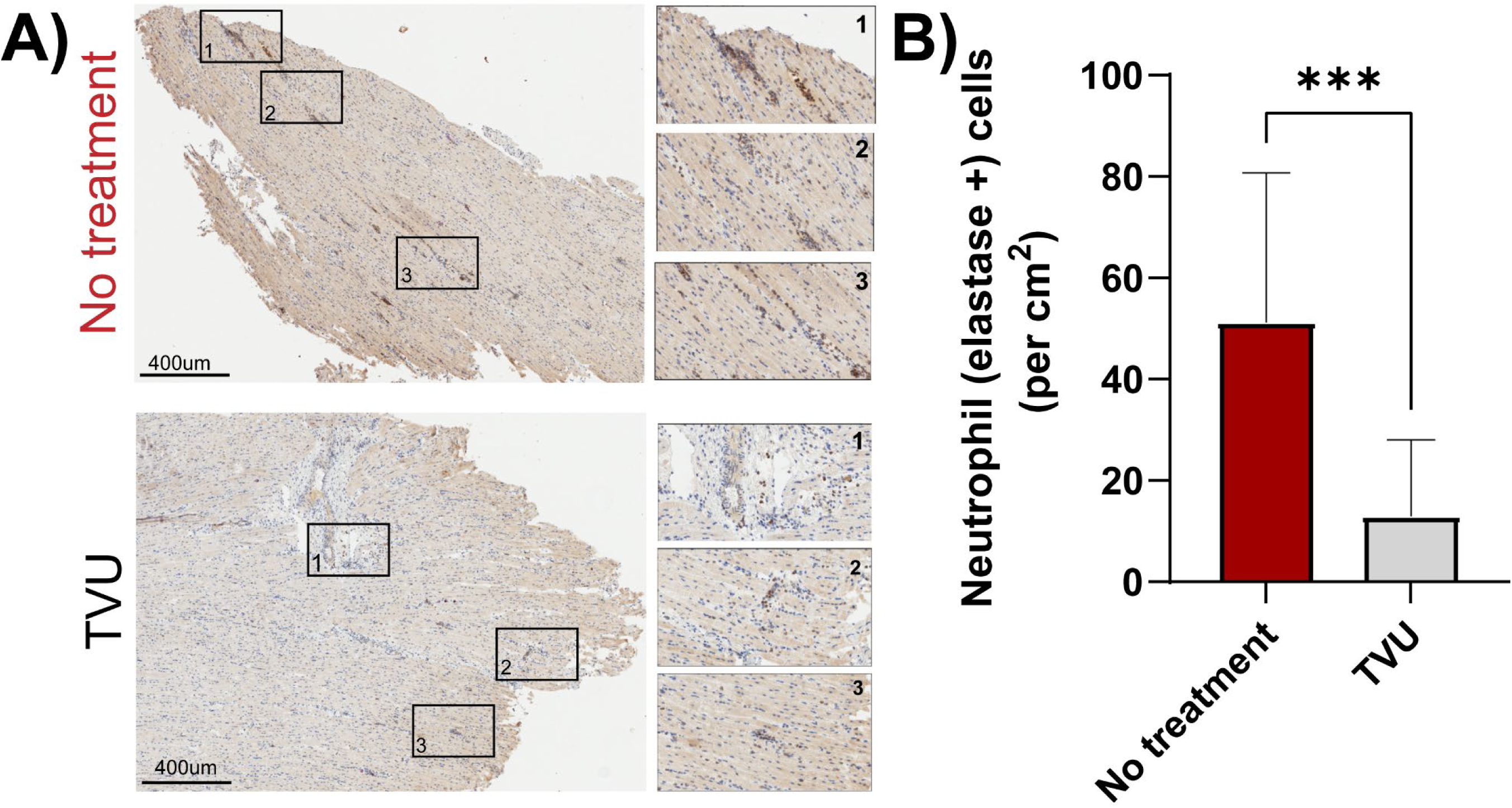
Histological observations of inflammation. **A)** Representative images and **B)** summary data showing extensive neutrophil infiltration within control hearts (no treatment; n = 3), which was markedly reduced in TVU hearts (n=5). Samples taken from transverse ring 1, *** p < 0.001 by Mann-Whitney test.

### TVU Ameliorates Maladaptive Remodeling at Intercalated Discs Following MI/R

Ventricular arrhythmias, including sustained and non-sustained VT and VF, are well-recognized complications of acute MI, occurring most commonly in the early peri-infarct and reperfusion period and contributing substantially to morbidity and in-hospital mortality^27–30^. The cardiomyocyte intercalated disc (ID) is a site of pro-arrhythmic remodeling following MI/R. Inflammation and edema - triggered disruption of gap junctions and mislocalization of Na_V_1.5 away from juxta-junctional ID nanodomains have been linked with increased arrhythmia vulnerability^15,25,31–36^. Therefore, we performed confocal microscopy to characterize the ID structure and examine patterns of Cx43, N-cad, and Na_V_1.5 localization.

Infarct regions sampled from control hearts post MI/R showed diffuse distribution of Cx43 within IDs in sharp contrast to similar regions from TVU hearts, which displayed the dense, discrete punctate organization observed in healthy myocardium (Fig. 6)^37^.We quantified Cx43 cluster volume from confocal images. We observed a significant increase in Cx43 cluster volumes in the control group compared to the TVU group within infarct regions, whereas no such remodeling was noted in remote myocardium in either group. Overall, these data are consistent with the disruption of gap junction plaques by IRI being ameliorated by TVU.

To test the integrity of Nav1.5 clustering in a setting of human MI/R, we performed histological studies on cardiac tissue samples obtained from patients with acute MI (Fig. 5). Tissue samples were collected from the infarct region and remote myocardium (millimeters away from the infarct) as illustrated in (Fig. 5). Within infarct regions, we observed a significant reduction in spatial co-expression of Na_V_1.5 with the mechanical junction protein, N-cadherin. This prompted us to investigate the localization of Na_V_1.5 channels relative to junctional proteins within intercalated discs in the aftermath of MI/R. In this context, the Cx43 remodeling in control hearts following MI/R as described above precluded the use of Cx43 as a structural fiducial for Na_V_1.5 localization. Therefore, we undertook confocal microscopy studies to examine Na_V_1.5 localization relative to N-cadherin (N-cad) in our porcine MI/R model. Whereas Na_V_1.5 is densely clustered near N-cad and Cx43 in healthy myocardium^38,39^, we observed diffuse localization of Na_V_1.5 within IDs from infarct-adjacent regions following MI/R in control hearts. Both infarct and remote myocardial regions showed diffuse distributions of Na_V_1.5 at the ID with poor association with N-cad in the untreated control group. Such remodeling was only evident in the infarct regions from TVU hearts (Fig. 7A). To quantify this effect, we applied our previously validated point process analysis-based spatial statistical approach, Spatial Pattern Analysis using Closest Events (SPACE; Fig. 7B). Briefly, the cumulative distribution of nearest neighbor distances between Na_V_1.5 and N-cad immunosignals was compared against predicted distributions under complete spatial randomness to calculate indices of non-random spatial association^22^. Non-random spatial association of Na_V_1.5 with N-cad was reduced in the remote myocardium from control hearts relative to TVU hearts. A similar effect was observed at infarct sites. However, it did not reach statistical significance (p = 0.09). Overall, these data are consistent with MI/R inducing Na_V_1.5 remodeling at intercalated discs and suggest that such potentially proarrhythmic impact is mitigated by TVU.

**Figure 5:**
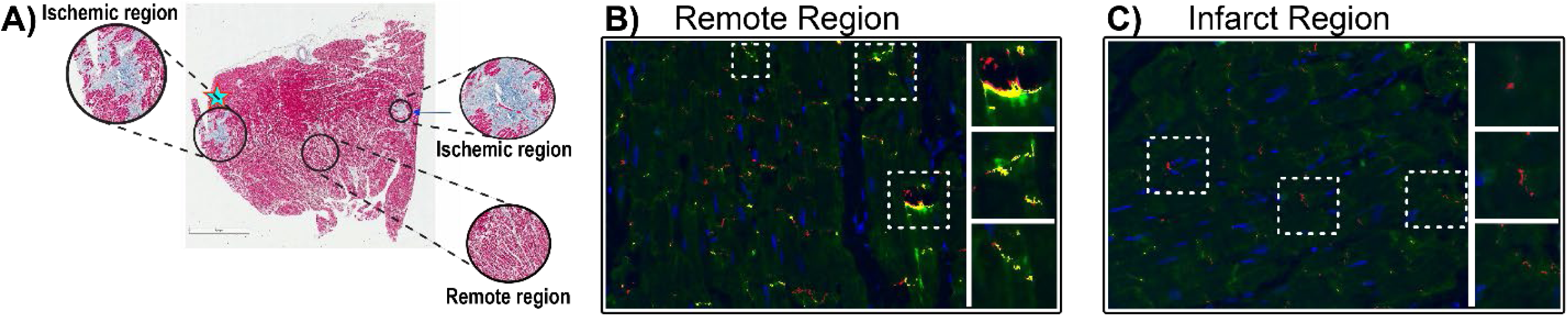
ID Remodeling in Acute MI patients. **A)** Representative H&E image from a patient with acute MI identifying infarcts and remote myocardium. **B)** Representative immunofluorescence images demonstrating robust co-distribution of Na_V_1.5 (green) and N-cad (red; co-localization indicated in yellow) at IDs in remote regions but not infarct regions.

**Figure 6:**
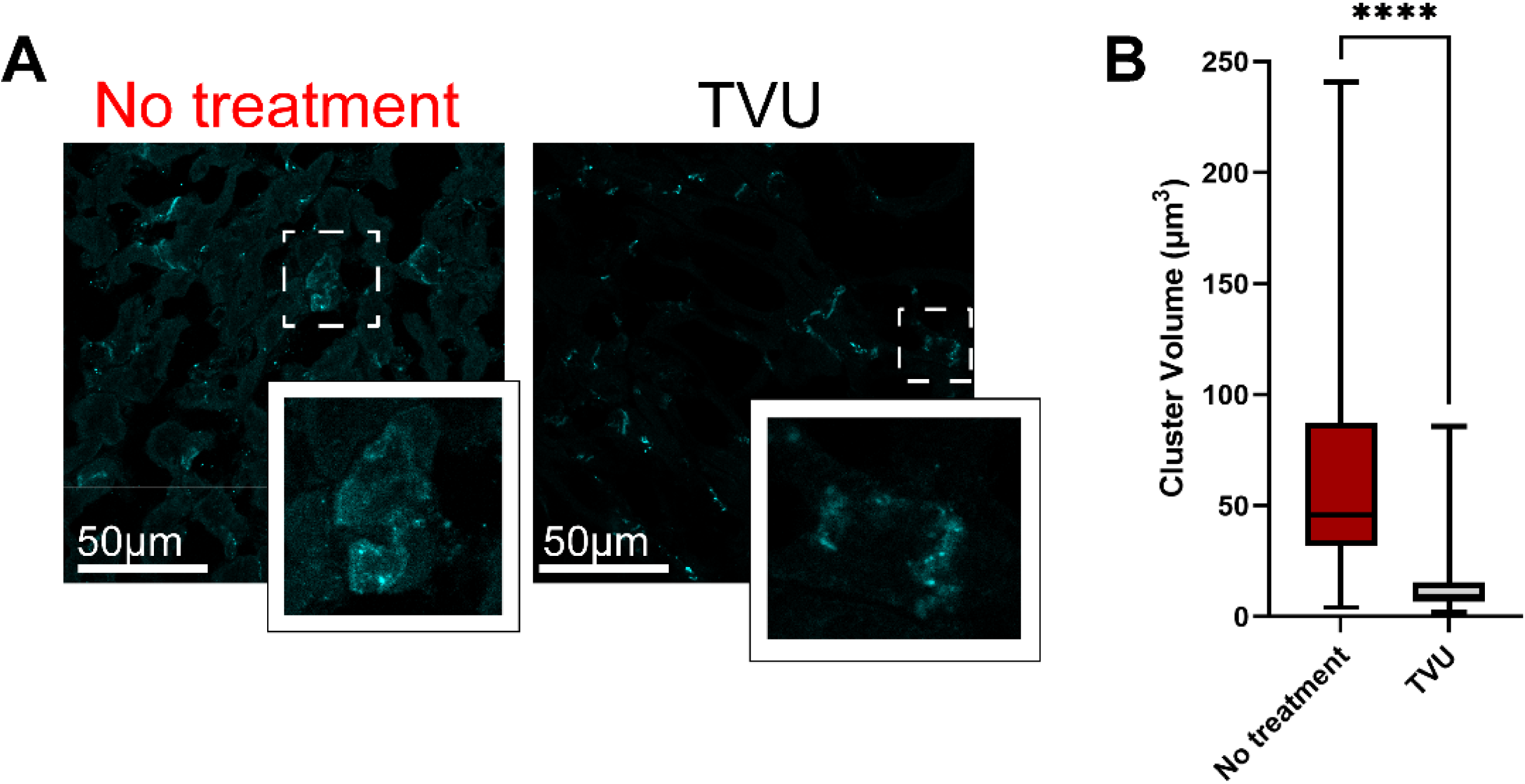
Cx43 remodeling post MI/R. A). Representative immunofluorescence images showing diffuse Cx43 signal within IDs from infarcted regions in control (No treatment) hearts, which is contrasted by clear, punctate Cx43 substructure observable at corresponding sites in TVU hearts. **B)** Summary plot of Cx43 cluster volume, 51 ID sites analyzed from n = 4 hearts in each group, **** p <0.0001 by Kolmogorov-Smirnov test.

**Figure 7:**
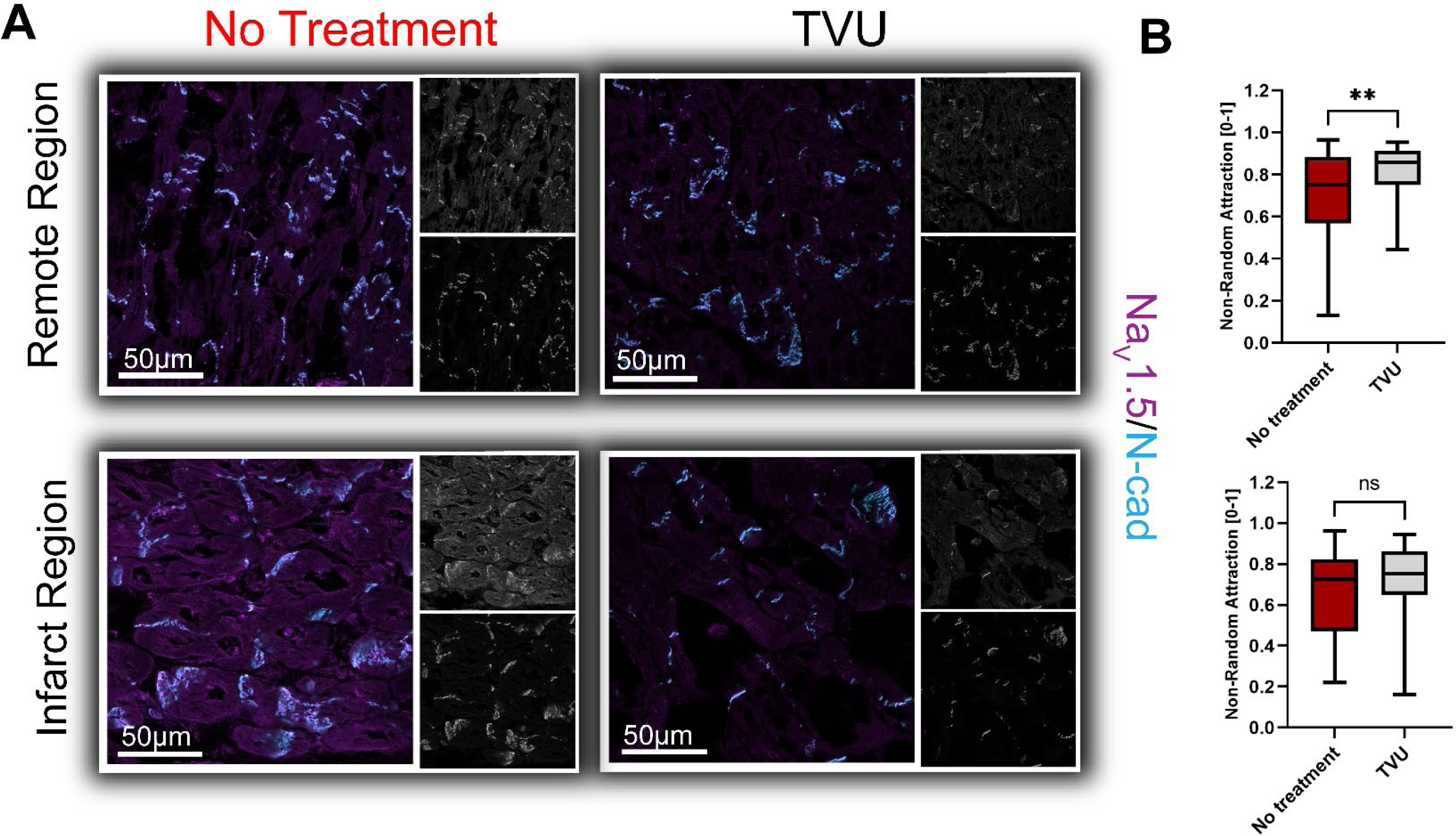
Mitigation of Na_V_1.5 remodeling by TVU. **A)** Representative immunofluorescence images showing N-cadherin (cyan) and Na_V_1.5 (magenta) localization within IDs from infarct and remote myocardium from our porcine MI/R studies. **B)**Summary plots of Na_V_1.5’s attraction to N-Cad. Remote region (TVU – 55 ID sites from 5 hearts, No treatment - 47 ID sites from 4 hearts); Infarct region (TVU – 61 ID sites from 5 hearts, No treatment – 42 ID sites from 4 hearts); ** p < 0.01, two-sample Kolmogorov-Smirnov test.

### TVU Mitigates VT inducibility following MI/R

Finally, we sought to assess the influence of TVU on MI/R-induced pro-arrhythmia by correlating the structural findings described above with functional observations collected through *in vivo* electrocardiography. ECG data were separated into experimental epochs corresponding to occlusion (90 minutes) and reperfusion (120 minutes) and durations of arrhythmic episodes compared between study groups. Although no difference in arrhythmia duration between TVU and untreated control hearts was apparent during the occlusion period, TVU hearts had significantly lower arrhythmia duration during the reperfusion phase (Fig. 8). Overall, all animals in both groups experienced episodes of VT with median durations of 3 ± 10.9 s and 2 ± 1.6 s in the control and TVU groups respectively (p < 0.01 between groups). This difference was reflective of a marked difference between the groups in the incidence of sustained VT. Control pigs experienced a median of 14 episodes (quartiles: 9, 18 episodes/animal; 6/6 pigs; mean duration: 16.8 ± 11 s) lasting 5 seconds or longer compared to 6 episodes in TVU pigs (quartiles: 2, 16 episodes/animal; 4/5 pigs; mean duration: 6.6 ± 0.7 s). Whereas all control pigs experienced VT episodes lasting 10 seconds or longer (median 5 episodes/animal; quartiles: 5, 6 episodes/animal; mean duration: 28.5 ± 17.6 s), only two of five pigs in the TVU experienced such episodes (median: 0 episodes/animal; quartiles: 0, 2 episodes/animal; mean duration: 11.9 ± 0.2 s). Across both study groups, the number and duration of VT episodes longer than 10 seconds were positively correlated with infarct area normalized to the area at risk with correlation coefficients of 0.71 and 0.58 respectively. These data collectively suggest that TVU significantly reduces arrhythmia burden following MI/R, especially the incidence and duration of sustained VTs.

**Figure 8:**
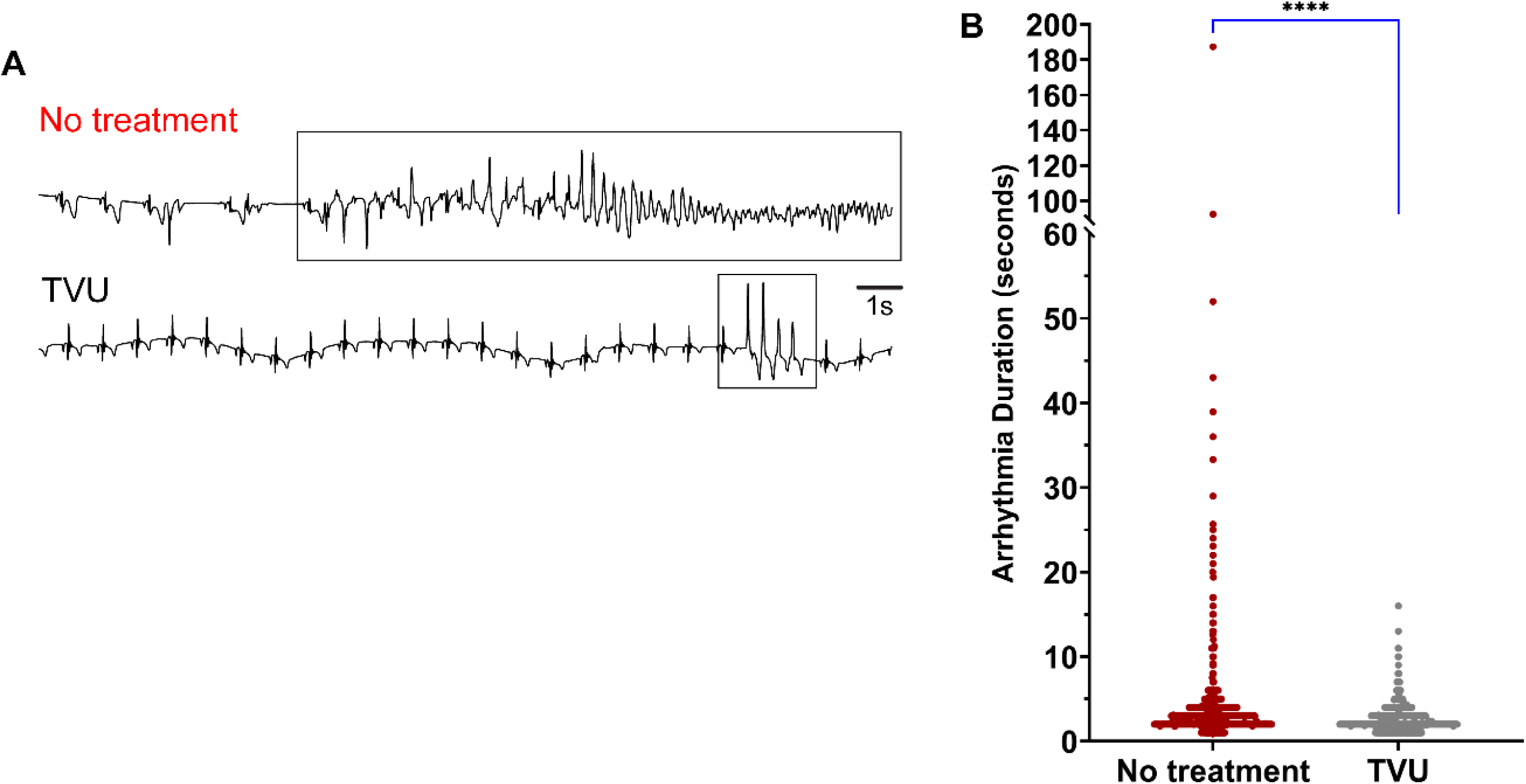
TVU mitigates sustained VT burden. **A)** Representative ECG traces from the reperfusion period illustrating sustained VT observed in a control pig (No treatment) contrasted by rapid termination of arrhythmia in a TVU pig. **B)** Summary plot of arrhythmia durations (TVU: n = 5 pigs, No treatment: n = 6 pigs; **** p <0.0001 by Kolmogorov-Smirnov test).

## DISCUSSION

In this study, we demonstrate that transvalvular unloading (TVU) before reperfusion reduces myocardial injury, limits microvascular obstruction, attenuates neutrophil infiltration into the heart, reduces perivascular & interstitial edema, preserves intercalated disc organization, and reduces ventricular arrhythmia burden in a porcine model of acute myocardial ischemia/reperfusion (MI/R). Importantly, the antiarrhythmic effect of TVU was most evident during reperfusion, when untreated control animals developed longer and more frequent sustained ventricular tachycardia episodes. These functional observations were accompanied by preservation of connexin43 organization and stronger spatial association between Na_V_1.5 and N-cadherin TVU animals, suggesting that TVU limits the acute formation of an arrhythmogenic substrate by protecting both the coronary microcirculation and the nanoscale architecture required for coordinated impulse propagation. Together, these findings support a mechanistic model in which unloading prior to reperfusion interrupts the cascade linking ischemia-reperfusion injury, microvascular failure, inflammatory edema, intercalated disc remodeling, and ventricular tachyarrhythmias.

As many as 300,000 patients undergo percutaneous coronary intervention (PCI) in the United States to treat acute myocardial infarction (AMI)^1,2^. Given the extensive body of evidence suggesting that maximizing myocardial salvage is key to improving patient outcomes following AMI, current clinical practice aims to restore perfusion as speedily as possible through PCI^3,4^. Paradoxically, however, reperfusion initially exacerbates myocardial injury, a phenomenon labeled myocardial ischemia-reperfusion injury (IRI)^5^, and contributes to increased arrhythmia burden, risk of heart failure, and other adverse outcomes in patients. The protective effects observed here extend prior work showing that left ventricular unloading before reperfusion can reduce ischemia-reperfusion injury and improve myocardial salvage^12,13^. In addition to confirming smaller infarcts, our data suggest that TVU modifies the tissue environment in which reperfusion injury evolves. By reducing ventricular load before flow restoration, TVU lowers wall stress and oxygen demand as previously demonstrated^40,41^, while improving the balance between epicardial reperfusion and microvascular perfusion. This distinction is important because successful opening of the culprit artery does not ensure effective tissue-level reperfusion. Persistent microvascular obstruction remains a major determinant of infarct expansion, adverse remodeling, and downstream heart failure risk^42–45^.

The marked reduction in MVO, neutrophil infiltration, and perivascular edema in TVU-treated hearts points to preservation of the microvascular circulation as a central component of this cardioprotection. MVO is caused by convergence of endothelial swelling, capillary plugging, microvascular spasm, thromboinflammation, and extravasation, all of which impair tissue perfusion despite restoration of epicardial flow^43,46^. In this context, the reduced inflammatory burden observed with TVU is unlikely to be secondary to smaller infarct size alone; rather, it may represent an upstream mechanism by which unloading stabilizes the reperfused microcirculation. This interpretation is consistent with the concept that microvascular injury and inflammation reciprocally amplify one another during reperfusion^47,48^ creating a local environment that promotes edema, conduction slowing, and electrical instability. Thus, TVU may provide benefit not only by salvaging myocardium, but also by limiting the vascular and inflammatory signals that convert reperfused tissue into a dynamic arrhythmogenic substrate.

A key implication of this work is that the antiarrhythmic effect of TVU may arise from preservation of cardiac structure at multiple spatial scales. At the tissue level, reduced infarct size and MVO would be expected to lessen spatial heterogeneity in excitability and conduction. At the cellular and subcellular levels, the preservation of connexin43 organization and Na_V_1.5 association with N-cadherin suggests that TVU protects intercalated disc nanodomains that coordinate electrical coupling and sodium current availability at sites of cell-cell contact. This is mechanistically important because acute inflammation and interstitial edema can disrupt intercalated disc architecture, disperse Na_V_1.5-rich nanodomains, and promote conduction defects that favor reentry^49,50^. The observation that human acute MI tissue also showed reduced Na_V_1.5/N-cadherin co-distribution near infarcted myocardium strengthens the translational relevance of this pathway and supports the idea that the porcine MI/R model meaningfully recapitulates clinically observed forms of arrhythmogenic remodeling. In this framework, TVU does not function as a conventional antiarrhythmic intervention that directly targets ion channel function or excitability; instead, its effects may be mediated by upstream mitigation of vascular and inflammatory injury that destabilize the structural underpinnings conduction during reperfusion.

These findings may help explain why pharmacologic strategies aimed at limiting reperfusion injury, inflammation, or gap junction remodeling have not consistently translated into clinical benefit. Many such interventions are administered at or after reperfusion, when microvascular damage, inflammatory activation, and electrical remodeling may already be underway. In contrast, TVU is applied before reperfusion and may therefore act earlier in the injury cascade, creating a more favorable hemodynamic and microvascular milieu before oxygenated blood re-enters ischemic myocardium. This temporal advantage may be particularly relevant for arrhythmia prevention, because reperfusion-associated ventricular arrhythmias often emerge rapidly and may be driven by transient but severe changes in perfusion, edema, excitability, and conduction. The present data therefore broaden the rationale for unloading beyond infarct-size reduction and suggest arrhythmia mitigation as an important mechanistic and translational endpoint to consider in future studies of unloading strategies.

### Limitations

We acknowledge a multitude of limitations, which should be recognized while interpreting our results. Foremost is that our MI/R studies were performed in healthy pigs lacking any of the comorbidities typically exhibited by patients experiencing AMI. Thus, future work in chronic disease models is needed. In order for the MI/R studies to be completed, pigs were administered the beta blocker metoprolol to stabilize them, which may have helped mitigate MVO and other sequelae of MI/R. Nevertheless, we observed a marked difference in MVO and myocardial health between our study groups despite both receiving similar doses of metoprolol, suggesting that TVU provided significant therapeutic benefits. The spectral footprint in thioflavin-S necessitated the use of fluorescence lifetime resolved confocal microscopy and precluded super-resolution light microscopy methods to assess protein localization with nanoscale resolution. Likewise, the time course of imaging thioflavin-S precluded immediate preservation of tissue samples for electron microscopy. Future studies are needed using such techniques to directly assess the impact of TVU on cardiac nanostructure. Humoral and cellular inflammation are complex processes with many downstream factors, which merit in-depth analysis in future studies.

## Conclusions

In summary, we demonstrate using a porcine preclinical model that that transvalvular unloading applied prior to the onset of reperfusion improves myocardial salvage and mitigates arrhythmia burden following acute myocardial infarction. Further, we provide evidence that these benefits derive from mitigation of inflammation and preservation of microvascular integrity, effects which could positively impact patients’ downstream risk of heart failure. We also demonstrate how TVU may help preserve normal action potential conduction in the heart by protecting intercalated disc nanodomains, which are disrupted in arrhythmia patients including those with AMI. Thus, our work highlights TVU as a therapeutic option with significant clinical potential for mitigating arrhythmias following AMI.

## Availability of data and material

Raw experimental data are backed up on UNR servers and will be shared freely on request.

## Code availability

Image analysis was performed using open-source software previously developed and validated by the corresponding author’s group. Source code is freely available via Github.

## REFERENCES

1. Afana M, Brinjikji W, Cloft H, Salka S. Hospitalization costs for acute myocardial infarction patients treated with percutaneous coronary intervention in the United States are substantially higher than Medicare payments. Clin Cardiol. 2015;38:13–19. doi: 10.1002/clc.22341

2. Cram P, House JA, Messenger JC, Piana RN, Horwitz PA, Spertus JA. Indications for percutaneous coronary interventions performed in US hospitals: a report from the NCDR(R). American heart journal. 2012;163:214–221 e211. doi: 10.1016/j.ahj.2011.08.024

3. Rao SV, O’Donoghue ML, Ruel M, Rab T, Tamis-Holland JE, Alexander JH, Baber U, Baker H, Cohen MG, Cruz-Ruiz M, et al. 2025 ACC/AHA/ACEP/NAEMSP/SCAI Guideline for the Management of Patients With Acute Coronary Syndromes: A Report of the American College of Cardiology/American Heart Association Joint Committee on Clinical Practice Guidelines. Circulation. 2025;151:e771–e862. doi: 10.1161/CIR.0000000000001309

4. Saha T, Soliman-Aboumarie H. Review of Current Management of Myocardial Infarction. J Clin Med. 2025;14. doi: 10.3390/jcm14176241

5. Welt FGP, Batchelor W, Spears JR, Penna C, Pagliaro P, Ibanez B, Drakos SG, Dangas G, Kapur NK. Reperfusion Injury in Patients With Acute Myocardial Infarction: JACC Scientific Statement. Journal of the American College of Cardiology. 2024;83:2196–2213. doi: 10.1016/j.jacc.2024.02.056

6. Curran J, Burkhoff D, Kloner RA. Beyond Reperfusion: Acute Ventricular Unloading and Cardioprotection During Myocardial Infarction. J Cardiovasc Transl Res. 2019;12:95–106. doi: 10.1007/s12265-019-9863-z

7. Esposito ML, Zhang Y, Qiao X, Reyelt L, Paruchuri V, Schnitzler GR, Morine KJ, Annamalai SK, Bogins C, Natov PS, et al. Left Ventricular Unloading Before Reperfusion Promotes Functional Recovery After Acute Myocardial Infarction. Journal of the American College of Cardiology. 2018;72:501–514. doi: 10.1016/j.jacc.2018.05.034

8. Fudim M, Konecny F, Heuring JJ, Durst CA, Fain ES, Patel MR. Left Ventricular Unloading Using Intra-aortic Entrainment Pumping Before Reperfusion Reduces Post-AMI Infarct Size. Journal of cardiac failure. 2025;31:538–548. doi: 10.1016/j.cardfail.2024.07.022

9. Hansen ESS, Madsen TL, Wood G, Granfeldt A, Bogh N, Tofig BJ, Agger P, Lindhardt JL, Poulsen CB, Botker HE, et al. Veno-occlusive unloading of the heart reduces infarct size in experimental ischemia-reperfusion. Sci Rep. 2021;11:4483. doi: 10.1038/s41598-021-84025-y

10. Kapur NK, Reyelt L, Swain L, Esposito M, Qiao X, Annamalai S, Meyns B, Smalling R. Mechanical Left Ventricular Unloading to Reduce Infarct Size During Acute Myocardial Infarction: Insight from Preclinical and Clinical Studies. J Cardiovasc Transl Res. 2019;12:87–94. doi: 10.1007/s12265-019-09876-3

11. Swain L, Reyelt L, Bhave S, Qiao X, Thomas CJ, Zweck E, Crowley P, Boggins C, Esposito M, Chin M, et al. Transvalvular Ventricular Unloading Before Reperfusion in Acute Myocardial Infarction. Journal of the American College of Cardiology. 2020;76:684–699. doi: 10.1016/j.jacc.2020.06.031

12. Saku K, Kakino T, Arimura T, Sunagawa G, Nishikawa T, Sakamoto T, Kishi T, Tsutsui H, Sunagawa K. Left Ventricular Mechanical Unloading by Total Support of Impella in Myocardial Infarction Reduces Infarct Size, Preserves Left Ventricular Function, and Prevents Subsequent Heart Failure in Dogs. Circ Heart Fail. 2018;11:e004397. doi: 10.1161/CIRCHEARTFAILURE.117.004397

13. Sun X, Li J, Zhao W, Lu S, Guo C, Lai H, Wang C. Early Assistance With Left Ventricular Assist Device Limits Left Ventricular Remodeling After Acute Myocardial Infarction in a Swine Model. Artif Organs. 2016;40:243–251. doi: 10.1111/aor.12541

14. Mezache L, Nuovo GJ, Suster D, Tili E, Awad H, Radwański PB, Veeraraghavan R. Histologic, viral, and molecular correlates of heart disease in fatal COVID-19. Annals of Diagnostic Pathology. 2022:151983. doi: 10.1016/j.anndiagpath.2022.151983

15. Mezache L, Soltisz A, Tili E, Nuovo GJ, Veeraraghavan R. SARS-CoV-2 spike protein-induced inflammation underlies proarrhythmia in COVID-19. Sci Rep. 2025;15:33991. doi: 10.1038/s41598-025-12807-9

16. Magro C, Mulvey JJ, Berlin D, Nuovo G, Salvatore S, Harp J, Baxter-Stoltzfus A, Laurence J. Complement associated microvascular injury and thrombosis in the pathogenesis of severe COVID-19 infection: A report of five cases. Transl Res. 2020;220:1–13. doi: 10.1016/j.trsl.2020.04.007

17. Perez-Bermejo JA, Kang S, Rockwood SJ, Simoneau CR, Joy DA, Silva AC, Ramadoss GN, Flanigan WR, Fozouni P, Li H, et al. SARS-CoV-2 infection of human iPSC-derived cardiac cells reflects cytopathic features in hearts of patients with COVID-19. Sci Transl Med. 2021;13. doi: 10.1126/scitranslmed.abf7872

18. Varga Z, Flammer AJ, Steiger P, Haberecker M, Andermatt R, Zinkernagel AS, Mehra MR, Schuepbach RA, Ruschitzka F, Moch H. Endothelial cell infection and endotheliitis in COVID-19. Lancet. 2020;395:1417–1418. doi: 10.1016/S0140-6736(20)30937-5

19. Mezache L, Soltisz AM, Johnstone SR, Isakson BE, Veeraraghavan R. Vascular Endothelial Barrier Protection Prevents Atrial Fibrillation by Preserving Cardiac Nanostructure. JACC Clin Electrophysiol. 2023. doi: 10.1016/j.jacep.2023.10.013.

20. Struckman HL, Moise N, King DR, Soltisz A, Buxton A, Dunlap I, Chen Z, Radwanski PB, Weinberg SH, Veeraraghavan R. Unraveling Impacts of Chamber-Specific Differences in Intercalated Disc Ultrastructure and Molecular Organization on Cardiac Conduction. JACC Clin Electrophysiol. 2023. doi: 10.1016/j.jacep.2023.05.042

21. Bogdanov V, Soltisz AM, Moise N, Ivanova M, Andreev I, Sakuta G, Weinberg SH, Davis JP, Veeraraghavan R, Györke S. Distributed synthesis of sarcolemmal and sarcoplasmic reticulum membrane proteins in cardiac myocytes. Bas Res Cardiol. 2021;116:63. doi: 10.1007/s00395-021-00895-3

22. Soltisz AM, Craigmile PF, Veeraraghavan R. Spatial Pattern Analysis using Closest Events (SPACE)-A Nearest Neighbor Point Pattern Analysis Framework for Assessing Spatial Relationships from Digital Images. Microsc Microanal. 2024;30:306–317. doi: 10.1093/mam/ozae022

23. Bertoluci MC, Ce GV, da Silva AM, Wainstein MV, Boff W, Punales M. Endothelial dysfunction as a predictor of cardiovascular disease in type 1 diabetes. World J Diabetes. 2015;6:679–692. doi: 10.4239/wjd.v6.i5.679

24. de Zeeuw D, Parving HH, Henning RH. Microalbuminuria as an early marker for cardiovascular disease. J Am Soc Nephrol. 2006;17:2100–2105. doi: 10.1681/ASN.2006050517

25. Mezache L, Struckman HL, Greer-Short A, Baine S, Gyorke S, Radwanski PB, Hund TJ, Veeraraghavan R. Vascular endothelial growth factor promotes atrial arrhythmias by inducing acute intercalated disk remodeling. Sci Rep. 2020;10:20463. doi: 10.1038/s41598-020-77562-5

26. Montezano AC, Dulak-Lis M, Tsiropoulou S, Harvey A, Briones AM, Touyz RM. Oxidative stress and human hypertension: vascular mechanisms, biomarkers, and novel therapies. Can J Cardiol. 2015;31:631–641. doi: 10.1016/j.cjca.2015.02.008

27. Frampton J, Ortengren AR, Zeitler EP. Arrhythmias After Acute Myocardial Infarction. The Yale journal of biology and medicine. 2023;96:83–94. doi: 10.59249/LSWK8578

28. Rymer JA, Wegermann ZK, Wang TY, Li S, Smilowitz NR, Wilson BH, Jneid H, Tamis-Holland JE. Ventricular Arrhythmias After Primary Percutaneous Coronary Intervention for STEMI. JAMA Netw Open. 2024;7:e2410288. doi: 10.1001/jamanetworkopen.2024.10288

29. Tran HV, Ash AS, Gore JM, Darling CE, Kiefe CI, Goldberg RJ. Twenty-five year trends (1986-2011) in hospital incidence and case-fatality rates of ventricular tachycardia and ventricular fibrillation complicating acute myocardial infarction. American heart journal. 2019;208:1–10. doi: 10.1016/j.ahj.2018.10.007

30. Wu T, Wang Y, Liu H, Zheng Y, Feng Y. Clinical efficacy of amiodarone in prehospital emergency treatment of myocardial infarction: a systematic review and meta-analysis. J Cardiothorac Surg. 2025;21:40. doi: 10.1186/s13019-025-03773-4

31. Beardslee MA, Lerner DL, Tadros PN, Laing JG, Beyer EC, Yamada KA, Kleber AG, Schuessler RB, Saffitz JE. Dephosphorylation and intracellular redistribution of ventricular connexin43 during electrical uncoupling induced by ischemia. Circulation research. 2000;87:656–662. doi: 10.1161/01.res.87.8.656

32. George SA, Calhoun PJ, Gourdie RG, Smyth JW, Poelzing S. TNFalpha Modulates Cardiac Conduction by Altering Electrical Coupling between Myocytes. Front Physiol. 2017;8:334. doi: 10.3389/fphys.2017.00334

33. Mezache L, Soltisz AM, Johnstone SR, Isakson BE, Veeraraghavan R. Vascular Endothelial Barrier Protection Prevents Atrial Fibrillation by Preserving Cardiac Nanostructure. JACC Clin Electrophysiol. 2023;9:2444–2458. doi: 10.1016/j.jacep.2023.10.013

34. Peretto G, Rizzo S, Menegon A, Villatore A, Veeraraghavan R, Vignale D, Castoldi V, Perani L, Canu T, Panzeri MC, et al. Intercalated Disc Abnormalities Are Linked to Arrhythmias in Inflammatory Cardiomyopathy. JACC Clin Electrophysiol. 2025;11:1097–1110. doi: 10.1016/j.jacep.2025.02.025

35. Peters NS, Wit AL. Myocardial architecture and ventricular arrhythmogenesis. Circulation. 1998;97:1746–1754. doi: 10.1161/01.cir.97.17.1746

36. Wit AL, Peters NS. The role of gap junctions in the arrhythmias of ischemia and infarction. Heart rhythm: the official journal of the Heart Rhythm Society. 2012;9:308–311. doi: 10.1016/j.hrthm.2011.09.056

37. Meens MJ, Kwak BR, Duffy HS. Role of connexins and pannexins in cardiovascular physiology. Cell Mol Life Sci. 2015;72:2779–2792. doi: 10.1007/s00018-015-1959-2

38. Lin X, Liu N, Lu J, Zhang J, Anumonwo JM, Isom LL, Fishman GI, Delmar M. Subcellular heterogeneity of sodium current properties in adult cardiac ventricular myocytes. Heart rhythm: the official journal of the Heart Rhythm Society. 2011;8:1923–1930. doi: 10.1016/j.hrthm.2011.07.016

39. Shy D, Gillet L, Abriel H. Cardiac sodium channel NaV1.5 distribution in myocytes via interacting proteins: the multiple pool model. Biochim Biophys Acta. 2013;1833:886–894. doi: 10.1016/j.bbamcr.2012.10.026

40. Kapur NK, Qiao X, Paruchuri V, Morine KJ, Syed W, Dow S, Shah N, Pandian N, Karas RH. Mechanical Pre-Conditioning With Acute Circulatory Support Before Reperfusion Limits Infarct Size in Acute Myocardial Infarction. JACC Heart Fail. 2015;3:873–882. doi: 10.1016/j.jchf.2015.06.010

41. Meyns B, Stolinski J, Leunens V, Verbeken E, Flameng W. Left ventricular support by catheter-mounted axial flow pump reduces infarct size. Journal of the American College of Cardiology. 2003;41:1087–1095. doi: 10.1016/s0735-1097(03)00084-6

42. de Waha S, Patel MR, Granger CB, Ohman EM, Maehara A, Eitel I, Ben-Yehuda O, Jenkins P, Thiele H, Stone GW. Relationship between microvascular obstruction and adverse events following primary percutaneous coronary intervention for ST-segment elevation myocardial infarction: an individual patient data pooled analysis from seven randomized trials. European heart journal. 2017;38:3502–3510. doi: 10.1093/eurheartj/ehx414

43. Kloner RA, Ganote CE, Jennings RB. The “no-reflow” phenomenon after temporary coronary occlusion in the dog. The Journal of clinical investigation. 1974;54:1496–1508. doi: 10.1172/JCI107898

44. Krug A, Du Mesnil de R, Korb G. Blood supply of the myocardium after temporary coronary occlusion. Circulation research. 1966;19:57–62. doi: 10.1161/01.res.19.1.57

45. Orn S, Manhenke C, Greve OJ, Larsen AI, Bonarjee VV, Edvardsen T, Dickstein K. Microvascular obstruction is a major determinant of infarct healing and subsequent left ventricular remodelling following primary percutaneous coronary intervention. European heart journal. 2009;30:1978–1985. doi: 10.1093/eurheartj/ehp219

46. Mukherjee D, Moliterno DJ. Achieving tissue-level perfusion in the setting of acute myocardial infarction. The American journal of cardiology. 2000;85:39C–46C. doi: 10.1016/s0002-9149(00)00818-3

47. Kalogeris T, Baines CP, Krenz M, Korthuis RJ. Cell biology of ischemia/reperfusion injury. Int Rev Cell Mol Biol. 2012;298:229–317. doi: 10.1016/B978-0-12-394309-5.00006-7

48. Seal JB, Gewertz BL. Vascular dysfunction in ischemia-reperfusion injury. Ann Vasc Surg. 2005;19:572–584. doi: 10.1007/s10016-005-4616-7

49. Veeraraghavan R, Lin J, Hoeker GS, Keener JP, Gourdie RG, Poelzing S. Sodium channels in the Cx43 gap junction perinexus may constitute a cardiac ephapse: an experimental and modeling study. Pffugers Arch. 2015;467:2093–2105. doi: 10.1007/s00424-014-1675-z

50. Veeraraghavan R, Lin J, Keener JP, Gourdie R, Poelzing S. Potassium channels in the Cx43 gap junction perinexus modulate ephaptic coupling: an experimental and modeling study. Pffugers Arch. 2016;468:1651–1661. doi: 10.1007/s00424-016-1861-2

